# A pharmacological chaperone stabilizer rescues the expression of the vast majority of pathogenic variants in a G protein-coupled receptor

**DOI:** 10.1101/2024.11.28.625821

**Authors:** Taylor L. Mighell, Ben Lehner

## Abstract

Reduced protein stability is the most frequent mechanism by which rare missense variants cause disease. A promising therapeutic avenue for treating destabilizing variants is pharmacological chaperones (PCs, also known as correctors or stabilizers), small molecules that bind to and stabilize target proteins. PCs have been approved as clinical treatments for specific variants, but protein energetics suggest their effects might be much more general. Here, we test this hypothesis for the first time by comprehensively quantifying PC efficacy for all missense variants in a human disease gene, the vasopressin 2 receptor (V2R), a G-protein coupled receptor in which loss-of-function variants cause nephrogenic diabetes insipidus (NDI). Strikingly, treatment with a PC rescues the expression of nearly all destabilized variants, with non-rescued variants identifying the drug’s binding site. Our results provide proof-of-principle that a single small molecule can rescue destabilizing variants throughout a protein’s structure. The application of this principle to other proteins should allow the development of effective therapies for many genetic diseases.

## Introduction

Rare genetic diseases pose a formidable challenge for global health. While the number of affected individuals for any disease is a small percentage of the population, in total as many as 300 million people are affected by rare diseases^1^. Despite recent progress in computational methods^2^, the identification of causal pathogenic variants and the determination of molecular mechanisms remains an arduous challenge^3,4^. Furthermore, developing effective therapies for genetic diseases where only a small number of patients carry each causal variant is extremely challenging.

The most frequent mechanism by which missense variants cause rare diseases is reduced protein stability. Large-scale experimental^5^ and computational^6,7^ surveys estimate that 40-60% of pathogenic variants are explained by loss of stability. Compensating for this reduced stability therefore represents a potentially general strategy to treat rare diseases. Pharmacological chaperones (PCs) are typically small molecules whose binding increases the thermostability and subsequent steady state expression level of a target protein. A striking example of PC success is in the treatment of cystic fibrosis, where a combination treatment of two PCs and a channel potentiator offers effective treatment for patients with the most frequent Phe508del and some other variants^8^. PCs – also referred to as protein stabilizers or correctors – have also been developed for non-membrane proteins, including clinically approved PCs for lysosomal storage disease proteins^9^ and the amyloidogenic transthyretin protein^10^, as well as experimental PCs for the most frequently mutated tumor suppressor protein, p53^11,12^. To maximize the effectiveness of PC therapy though, it is necessary to identify the mechanism of all pathogenic variation for a given protein, as well as the response to PC. These data do not currently exist for any protein.

Depending on the mechanism of action, PCs could have high specificity, rescuing only subsets of pathogenic variants localized in particular regions of the protein^13,14^. Alternatively, PCs could have largely nonspecific stabilizing effects that offset the destabilization by most variants in a protein, wherever they are located^12^. Most human proteins are marginally stable, meaning that only small changes in folding energy are required to produce large changes in the abundance of folded protein^15^. Indeed in a collated set of 223 experimentally determined changes in Gibbs free energy of folding (ΔΔG) values for membrane protein mutations, 91% were less than 3 kcal/mol^16^. Similarly, massive experimental mutagenesis of soluble proteins has confirmed the vast majority of variants only cause small changes in folding energy^17^, as do known pathogenic variants^5^. Such small changes in fold stability could potentially be easily compensated for by small molecule binding, which can confer similarly sized changes in free energy as mutations^18^. Moreover, provided that free energies combine mostly additively^12,19,20^, the stabilization conferred by small molecule binding should be largely independent of where a small molecule binds and where a mutation is located, provided the compound specifically binds the native folded state.

The vasopressin 2 receptor (V2R) is a G-protein coupled receptor (GPCR) with an important role in water homeostasis in the kidneys^21^. The hormone arginine vasopressin (AVP) is the primary endogenous ligand of V2R, and AVP levels regulate the permeability of the collecting duct of the kidneys: increased levels of AVP lead to increased permeability and a higher level of water reabsorption. Upon AVP binding, V2R adopts an active conformation and couples primarily with G_αs_ containing heterotrimeric G-proteins, leading to intracellular signaling that results in translocation of aquaporin-2 water channels to the plasma membrane^22^. When the function of V2R is lost, water reabsorption is compromised, leading to nephrogenic diabetes insipidus (NDI)^23^. Individuals with NDI suffer from chronic dehydration that can lead to severe clinical outcomes, and treatment options exist only for managing symptoms^24^. Highlighting the sensitivity and importance of this system, rare gain-of-function mutations that increase the basal activity of V2R cause an opposite clinical designation of nephrogenic syndrome of inappropriate antidiuresis (NSIAD)^25^. The gene encoding V2R, *AVPR2*, is on the X-chromosome. The majority of NDI-affected individuals are hemizygous males, although skewed X-inactivation has been reported to cause NDI or subclinical phenotypes in some females^26^. Hundreds of *AVPR2* variants have been found in individuals with NDI, of which about half are missense variants and the remainder being nonsense, small insertions or deletions, or splice-site mutations^27^. Only a fraction of the missense variants have been experimentally characterized ^22,27–34^.

Here we use V2R as a model system to directly test the hypothesis that PCs can rescue destabilizing mutants across the structure of a complex human protein. First, we employ a multiplexed assay to quantify the effects of all possible variants on the cell surface expression of V2R, revealing that more than half the known pathogenic variants strongly impair V2R expression, as do thousands of other missense variants throughout the protein. Strikingly, treatment with the V2R small molecule binder Tolvaptan rescues the expression of nearly all these variants. Only a small number are not compensated for, with these identifying functionally important sites and the drug binding site. Our results provide proof that a single small molecule can rescue the expression of reduced expression mutations throughout the structure of a protein. The application of this principle to other proteins should allow the development of general stabilizers for many different genetic diseases.

## Results

### Massively parallel measurement of V2R surface expression

We used SUNi mutagenesis^35^ to generate a saturation variant library containing all single amino acid changes to the V2R coding sequence. Mutagenesis primers were designed to introduce a degenerate NNK or NNS codon at each position (depending on the wild-type codon, K=G or T, S=G or C) and random DNA barcodes were subsequently inserted into the plasmid backbone to enable identification of each variant with short-read sequencing. In order to link the full V2R variant sequence with the short DNA barcode, we used long read sequencing and successfully linked 66,031 barcode-variants (Supplemental Fig. 1a). In total, 7,005 out of a possible 7,400 missense and nonsense variants (94.7% of possible) were represented by at least one barcode, with a median of 5 barcodes per variant.

We implemented a fluorescence-activated cell sorting (FACS)-based approach to measure the surface expression of the library of variants in human cells^36,37^. First, the library of plasmids was recombined into HEK293T landing pad cells^38^ (Materials and Methods), ensuring that each cell carried exactly one variant. The V2R construct had a HA-epitope tag at the N-terminus, which would be extracellular in a properly folded and trafficked receptor. Therefore, we performed immunostaining with a fluorescent antibody but without permeabilizing the cells, so that only receptors that had reached the membrane would contribute to signal. Then we sorted cells into four bins, isolated DNA from the cells in each bin, and used short-read sequencing to count the frequency of each variant across the bins (Fig. 1a).

**Figure 1.**
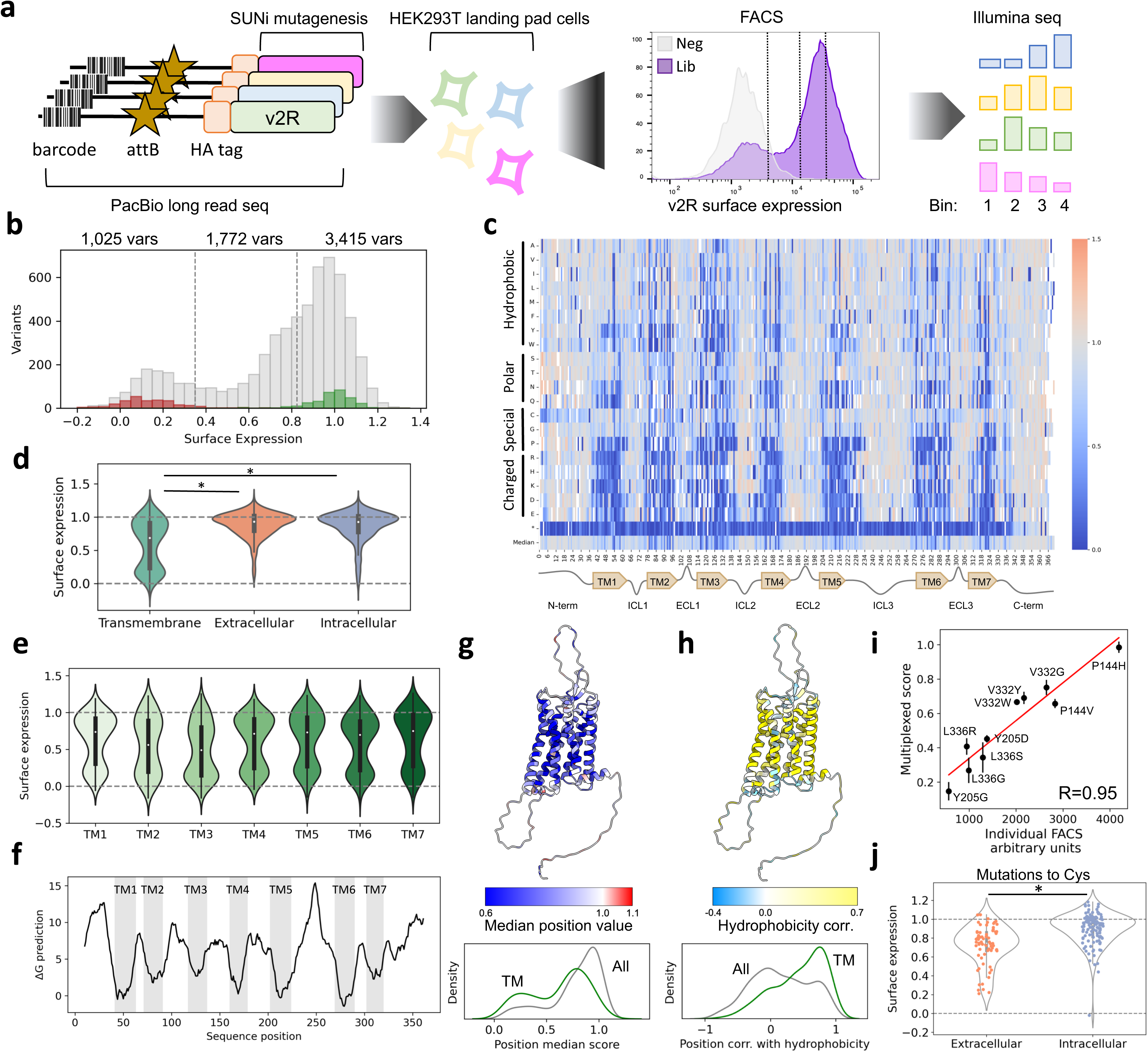
Experimental approach and primary dataset overview. **a** Experimental overview: a library of V2R variants was generated with SUNi mutagenesis and barcoded with random DNA barcodes. PacBio long-read sequencing was used to link variants with barcodes. Then, this library was integrated into HEK293T LLP-iCasp9-Blast landing pad cells. Fluorescent antibody staining followed by FACS was used to fractionate cells by expression level, then DNA was sequenced with Illumina short reads to identify the variant abundances in each bin. **b** Histogram of variant effect scores: premature truncating variants are in red, synonymous wildtype are in green, and missense variants are in gray. Dashed lines indicate the thresholds between well expressed, moderately expressed, and poorly expressed. The number of missense variants in each category is reported above. **c** Heatmap representation of the data. V2R receptor positions are on the x-axis, while mutant amino acid identities are on the y-axis. Blue colored cells indicate surface expression below wildtype; gray indicates same as wildtype; red indicates above wildtype expression. **d** Comparison of variant effects at positions within transmembrane helices, or in extra- or intra-cellular loops. **e** Comparison of variant effects at positions in transmembrane helices 1-7. **f** Predicted ΔG of membrane insertion (ΔG prediction server v1.0, Full protein scan, helix length=21). **g** Median position surface expression score colored on AlphaFold2 predicted V2R structure. Below: density plot of median position scores for the whole receptor (gray) or within transmembrane helices (green). **h** Per-position correlation of surface expression scores with Kyte-Doolittle hydrophobicity colored on AlphaFold2 predicted V2R structure. Below: density plot of per-position correlations for the whole receptor (gray) or within transmembrane helices (green). **i** Comparison of 10 variant scores measured individually or *en masse* (DMS measurements). Standard error of the mean shown for DMS measurements. Linear fit in red. **j** Effects of mutations to cysteine in the extra- or intra-cellular portions of the receptor.

Surface expression scores were calculated using the frequency of each variant in each bin multiplied by the geometric mean fluorescence value associated with each bin (Materials and Methods). Across four replicates we obtained high-confidence measurements for 6,844 (92.5% of possible, Materials and Methods) variants with high reproducibility (average pairwise replicate Pearson’s r=0.90, Supplementary Fig. 1b, data in Supplemental Table 1). Multiplexed scores correlate very well with FACS measurements of a series of isogenic cell lines bearing single V2R variants (r=0.95, Fig. 1i). Scores were normalized between complete loss of function (i.e. the median score of all nonsense variants in the first 300 positions of the receptor), assigned score 0, and the wild-type genotype assigned score 1. Synonymous wild-type variant scores are well separated from nonsense scores, and the distribution of all missense variants is bimodal, with a large fraction expressed near the level of wild-type and a smaller fraction expressed near the level of nonsense variants (Fig. 1b). We used the upper 95^th^ percentile of truncation scores (0.35) and the lower 95^th^ percentile of synonymous wild-type scores (0.825) to categorize missense variants as well expressed (3,415 variants), moderately expressed (1,772 variants), or poorly expressed (1,025 variants, Fig. 1b).

A heatmap representation of the data reveals that nonsense mutations are uniformly damaging through all seven transmembrane (TM) helices but mostly do not compromise surface expression in the unstructured, C-terminal tail (Fig. 1c). Transmembrane helices are particularly intolerant to mutation, especially to charged amino acids. In fact, mutations within the transmembrane helices as a population are significantly more damaging than the extra- or intracellular loops (Mann-Whitney U test, p=3.16x10^-135^, p=8.54x10^-153^, respectively), while there is no difference between the extra- and intra-cellular loops (p=0.39, Fig. 1d, g). Comparing the surface expression scores of all variants in each TM helix, variation in TM3 is the most damaging (median=0.49, Fig. 1e). This is consistent with previous work suggesting that TM3 has a critical role in receptor stability^39^ and acts as a “structural hub”^40^ for the TM bundle across class A GPCRs. TM2 is the second most mutation-sensitive (median=0.56) TM helix. The predicted free energy change (ΔG) of membrane integration^41^ for these helices is relatively unfavorable (Fig. 1f) suggesting that mutations here could further compromise an already inefficient process.

TM regions are solvated in lipid and are therefore enriched in hydrophobic residues, compared with extra- or intracellular residues. Mutations within the TM regions that decrease hydrophobicity have negative effects on surface expression (Spearman’s ρ=0.28, p=1.8x10^-48^) while the relationship is much weaker and in the opposite direction for the extra- or intracellular regions (ρ=-0.05, p=0.002, Supp. Fig. 1c). Visualization of each residue’s preference for hydrophobicity (calculated as rank correlation of surface expression scores with Kyte-Doolittle hydrophobicity) agrees with this notion (Fig. 1h) but also highlights several residues within the core of the receptor that prefer hydrophilic residues (Supplementary Fig. 1f).

V2R undergoes post-translational modification in the form of O- and N- glycosylation on the N-terminal tail^42^ as well as palmitoylation on the C-terminal tail.^43^ Positions 22-24 represent the N-glycosylation motif (N-X-S/T), and mutations at position 22 and 24 are much more deleterious than their neighbors in the unstructured N-terminus (Supplementary Fig. 1d). While O-glycosylation was reported at several serines and threonines in the N-terminus^42^, we only see strong mutational effects at S5 and T6, suggesting these are the only critical glycosylation sites, at least in HEK cells. Finally, palmitoylation at cysteines 341 and 342 have been reported to be important for surface expression^43^; our data show that in HEK cells mutations at these sites are well tolerated (Supplementary Fig. 1e).

### The contribution of V2R surface expression to pathogenicity

We next sought to understand the contribution of surface expression defects to V2R-related disease. To do this, we collated clinical variants from ClinVar^44^, population variants from gnomAD,^28^ and variants reported in individuals with NDI or NSIAD from the Human Gene Mutation Database^45^. There are striking differences between the surface expression scores for putatively benign variants (in gnomAD, or benign or likely benign in ClinVar) and the putatively loss-of-function ones (pathogenic or likely pathogenic in Clinvar, or NDI, Fig. 2a). There are no poorly expressed variants in the gnomAD, benign, or likely benign sets, and only 30.7% and 26.6% moderately expressed in likely benign and gnomAD, respectively (Fig. 2b). In contrast, 40%, 53.3%, and 55.6% of pathogenic, likely pathogenic, and NDI variants are poorly expressed, while 26.6%, 26.6%, and 30% are moderately expressed, respectively. There are only five known NSIAD variants and they are constitutively active^46^. Four are moderately and one is well expressed, consistent with the gain-of-function mechanism. Among variants of uncertain significance, 15.6% and 25% are poorly and moderately expressed, respectively (Fig. 2b). Stratifying gnomAD variants by allele frequency shows that common variants (allele frequency >0.001) are all well expressed (Fig. 2c).

**Figure 2.**
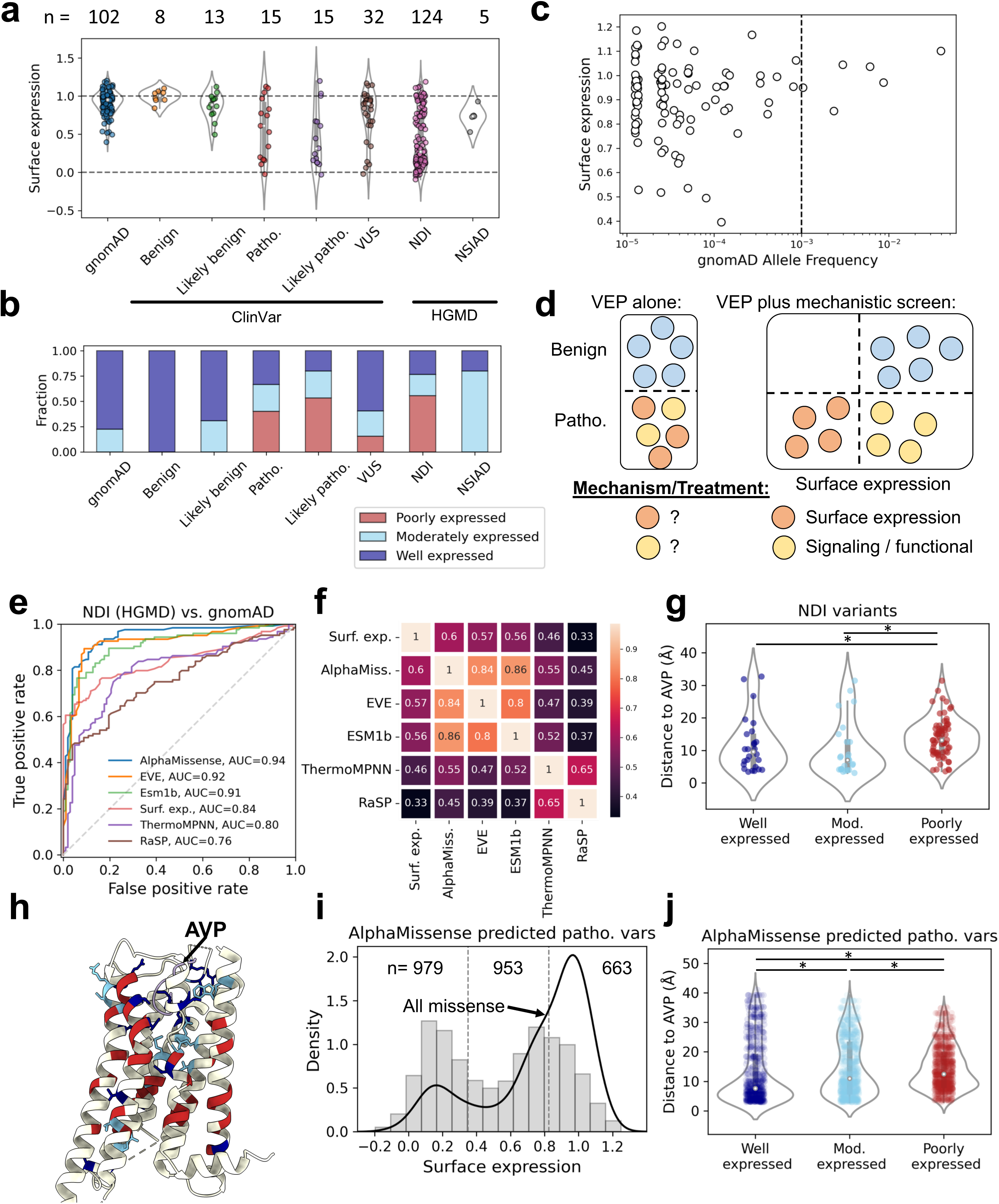
V2R clinical genetics and variant effect predictors. **a** Surface expression scores of V2R variants in gnomAD, ClinVar categories, or Human Gene Mutation Database (HGMD) categories. **b** Fraction of each variant category that is poorly-, moderately-, or well-expressed. **c** Relationship between allele frequency in gnomAD and surface expression score. Dashed line indicates allele frequency of 0.001, a commonly used threshold for common versus rare variation. **d** Schematic illustrating the importance of mechanistic screens. While VEPs are quite accurate for predicting pathogenicity, they cannot inform on the molecular mechanism of pathogenicity. **e** Receiver operating characteristic curve for different VEPS, or the empirical surface expression scores, for distinguishing NDI alleles from gnomAD alleles. **f** Correlations between the predictors and the empirical surface expression scores. **g** Distance between the endogenous ligand (AVP) and NDI variants of different expression levels. **h**

### Classifying pathogenic variant mechanisms

While computational variant effect predictors (VEPs) are increasingly accurate at distinguishing pathogenic from benign variation, they cannot perform the crucial task of determining molecular mechanisms for pathogenicity (Fig. 2d). First, we evaluated how well computational VEPs can discriminate pathogenic (NDI alleles) from putatively benign variation (gnomAD alleles). ESM1b^47^, a protein language model, achieves an area under the receiver operator characteristic curve (AUROC) of 0.91. EVE^48^, which uses multiple sequence alignments to model evolution, achieves AUROC of 0.92. Finally, AlphaMissense^49^, which uses structural predictions, population variant frequencies, and evolutionary data, achieves an AUROC of 0.94. As a comparison, surface expression scores achieve AUROC of 0.84, again suggesting that surface expression is a major determinant of pathogenicity.

On the other hand, models designed to predict protein stability effects of variation performed less well than the empirical surface expression scores. RaSP^50^ achieved AUROC of 0.76 and ThermoMPNN^51^ achieved 0.80 (Fig. 2e). We also compared the predictors to the surface expression scores and to each other. The VEPs correlated better with the surface expression data, with ρ=0.6, 0.57, and 0.56 for AlphaMissense, EVE, and ESM1b, respectively (Fig. 2f, Supplemental Fig. 2a-c). ThermoMPNN and RaSP correlated less well, at 0.46 and 0.33, respectively (Fig. 2f, Supplemental Fig. d-e). The poorer correlation of the stability predictors likely reflects the difficulty of predicting stability changes for membrane proteins, especially as RaSP and ThermoMPNN were trained on soluble proteins. Finally, we were curious whether mutation effect predictors could capture the pathogenicity of the NSIAD gain-of-function mutations. The number of variants is too small to make strong conclusions, but there seems to be much variability for the different models’ predictions, suggesting that gain-of-function variants could be more difficult to predict than loss-of-function (Supplemental Fig. 2a-e).

We hypothesized that our surface expression scores could help disambiguate the mechanism for pathogenic variants; specifically, whether a V2R variant is pathogenic due to decreased surface expression or a presumed signaling deficit. We considered that variation at positions close to the natural ligand (AVP) binding site in 3D space would be likelier to disrupt signaling than variation far away. Indeed, we found that well and moderately expressed NDI variants are closer to AVP in a solved structure (Mann-Whitney U test, p=1.5x10^-3^ and 7.7x10^-4^ for well and moderately expressed compared with poorly expressed variants, PDB structure 7KH0, Fig. 2g, h).

While there are over a hundred known NDI variants, this likely represents only a small fraction of pathogenic variation in V2R. Therefore, we used AlphaMissense to predict all pathogenic variants in V2R (AlphaMissense threshold=0.564^49^). 2,911 variants are predicted to be pathogenic, of which we have high confidence surface expression scores for 2,595. Of these, 979 (37.7%) are poorly expressed, 953 (36.7%) are moderately expressed, and 663 (25.5%) are well expressed (Fig. 2i). AlphaMissense predicted pathogenic variants have bimodal surface expression scores, just like the distribution of all missense variants. However, while the mode of the higher expression peak for all missense is near 1.0, the mode of the higher expression peak for AlphaMissense pathogenic variants is lower, around 0.8; this is likely due to a majority of these variants being in transmembrane helices (1,789 out of 2,595, 68.9%). The mode of the higher expression peak for transmembrane variants is also near 0.8 (see Fig. 1d). Finally, we compared the proximity of AlphaMissense pathogenic variants to AVP in the solved structure of V2R. As with the NDI variants, the well and moderately expressed pathogenic variants are closer to AVP in 3D space compared with the poorly expressed variants (p=2.2x10^-21^ and 0.03, respectively, Mann-Whitney U test, Fig. 2j), consistent with a specific signaling defect for variants that have at least moderate expression level. For this much larger set, the well expressed variants are also closer to AVP than the moderately expressed ones (p=1.9x10^-9^, Fig. 2j).

### Thermodynamic rescue of V2R variants

In principle, variants that are poorly expressed due to decreased thermodynamic stability should be rescued by incubating the cells expressing the variant library at reduced temperature^14^ (Fig. 3a). We sought to understand what fraction of V2R variants could be rescued by thermodynamic stabilization by culturing the cells expressing the V2R variant library at 27°. After 24 hours at 27°, FACS analysis revealed a shift in the distribution of surface expression, with more cells highly expressing V2R (Supplemental Fig. 3a). To gather quantitative data for all variants, we then sorted and sequenced the 27° rescued libraries. High confidence measurements were collected for 6,787 missense and nonsense variants (91.7% of possible) and replicates were well correlated (r=0.91, Supplemental Fig. 3c, data in Supplemental Table 1). The distribution of multiplexed missense variant scores mirrors the FACS data, with a shift of some variants from the low expressing to the high expressing peak (Fig. 3b).

**Figure 3.**
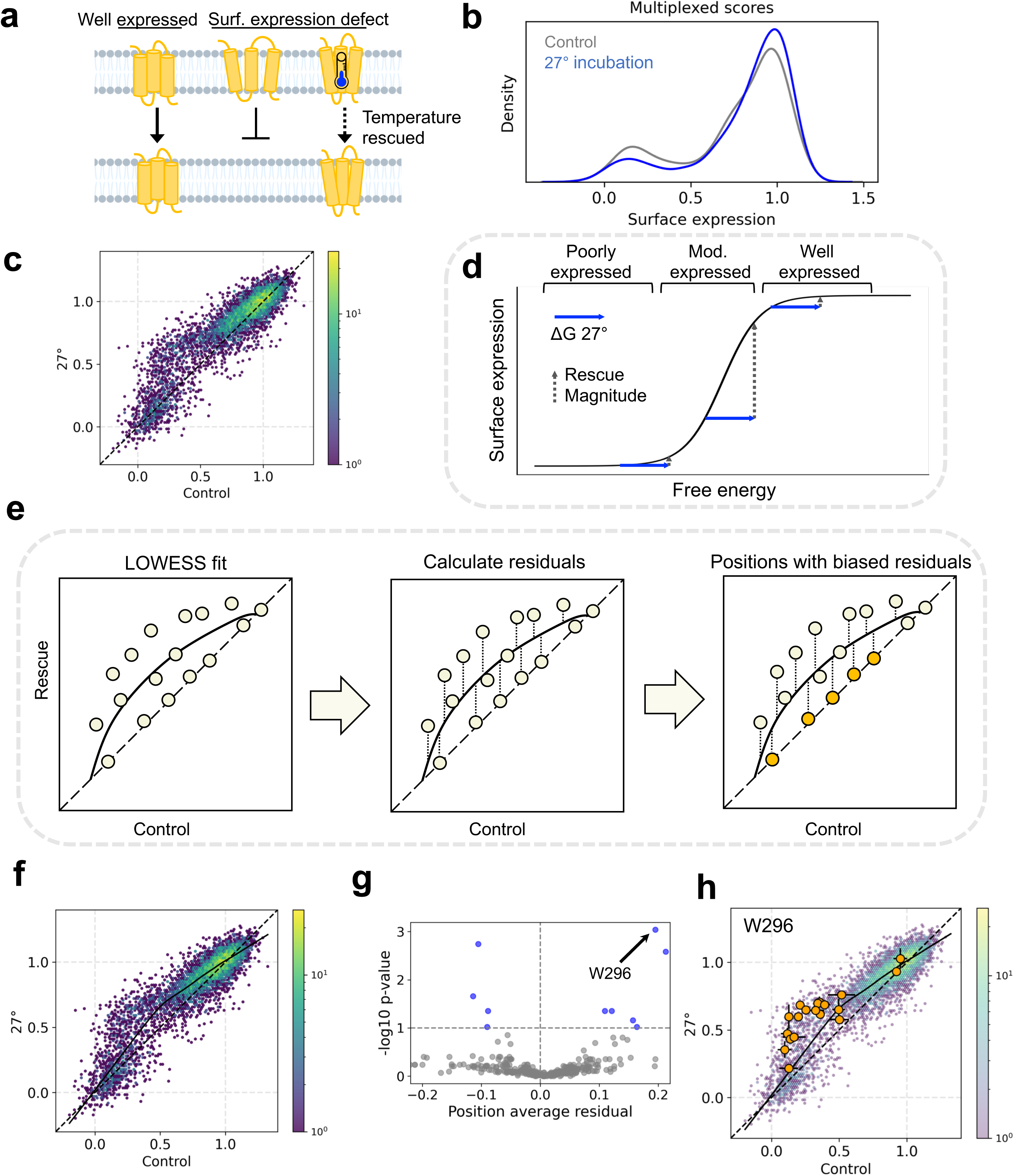
Thermodynamic rescue of V2R variants. **a** Schematic showing that well-folded receptors are properly trafficked to the membrane, but folding defective variants are not properly trafficked. These experiments seek to test which variants can be rescued by thermodynamic rescue. **b** Surface expression multiplexed score distribution of all missense variants in control (37°) and rescue (27° conditions). **c** Comparison of all missense variants in control and rescue condition. **d** Schematic representation of the relationship between free energy change and surface expression. The relationship between these two values is sigmoidal, meaning that the same free energy change (for example, that conferred by reducing culture temperature from 37° to 27°) can have different magnitude effects on surface expression. **e** The strategy employed to identify sites with greater or less rescue than expected. A Lowess curve is fit to the control versus rescue data; residuals to this line are calculated for all variants; positions are identified that have biased residuals. **f** Control versus rescue surface expression; Lowess fit in solid black; x=y line in dashed black. **g** Volcano plot comparing the average residual of all positions with the -log10 p-value. The position with the most significantly different set of residuals is highlighted (W296). **h** All missense variant effects at W296 are plotted with the whole population of missense variants in the background.

Then, we compared the surface expression of all variants at 27° compared to the control condition, 37°. The magnitude of rescue is greatest for variants with moderate expression level (Fig. 3c). On the other hand, variants that are well- or poorly-expressed in the control condition do not show large changes. This result is consistent with temperature reduction contributing a constant, additive free energy change, but that free energy change having a non-linear effect on the phenotype of interest^15,52^, in this case steady state expression level. Namely, the magnitude of rescue is greatest for variants with starting free energy values in the steepest part of the free energy-expression level curve (Fig. 3d).

To further investigate the generality of rescue afforded by temperature reduction, we designed an approach to identify positions that have more or less rescue compared to all other positions. First, we fit a Lowess curve to control vs. rescue data, which represents a null expectation for the magnitude of rescue at each control surface expression value. After fitting the curve, we calculate residuals between each variant and the fit line. Then, the residuals for each position are compared to the residuals at all other positions to find outliers (Fig. 3e). Namely, after fitting the Lowess curve, we used a two-sided Mann-Whitney U test to identify positions with significantly biased residuals (Materials and Methods, Supplemental Table 2). For the 27° condition, out of 371 positions, only four positions are rescued less than expected, while six are rescued more than expected for their expression level, (FDR=0.1, Fig. 3g-h). Among these ten positions are three glycosylation sites (positions 5, 6, and 24), which would be expected to have effects beyond simple thermodynamic destabilization. This suggests that the majority of variants can be rescued by thermodynamic stabilization.

### Pharmacological chaperone rescue of V2R variants

As thermodynamic stabilization demonstrated a generic rescue of the majority of V2R variants, we predicted that PC binding could also have a general rescue effect. PCs are typically hydrophobic small molecules that cross the plasma membrane and stabilize a target protein by binding to the folded state, so promoting trafficking (Fig. 4a). While PCs have shown effectiveness for correcting NDI-associated V2R variants both *in vitro* and in the clinic^53^, a major obstacle has been the uncertainty of how general the effect of any PC would be^22^. It could be that PCs have quite specific effects, such that each PC only rescues a subset of structurally-related variants^13,14^. Alternatively, PCs may generally rescue mutants with reduced thermodynamic stability^12^. Tolvaptan (also known as OPC-41061) is a V2R-specific, competitive, small molecule antagonist^54^ (Fig. 4b), and is approved for treating autosomal dominant polycystic kidney disease^55^, in which AVP-V2R signaling is misregulated. However, Tolvaptan has also been explored *in vitro* for its activity as a PC. In most tested cases, Tolvaptan not only rescues surface expression, it also enables some level of AVP-mediated signaling^32,34,56,57^. However, as for PCs in general, Tolvaptan has only been tested for effectiveness on a very limited set of variants (<15)^32,34,56,57^.

**Figure 4.**
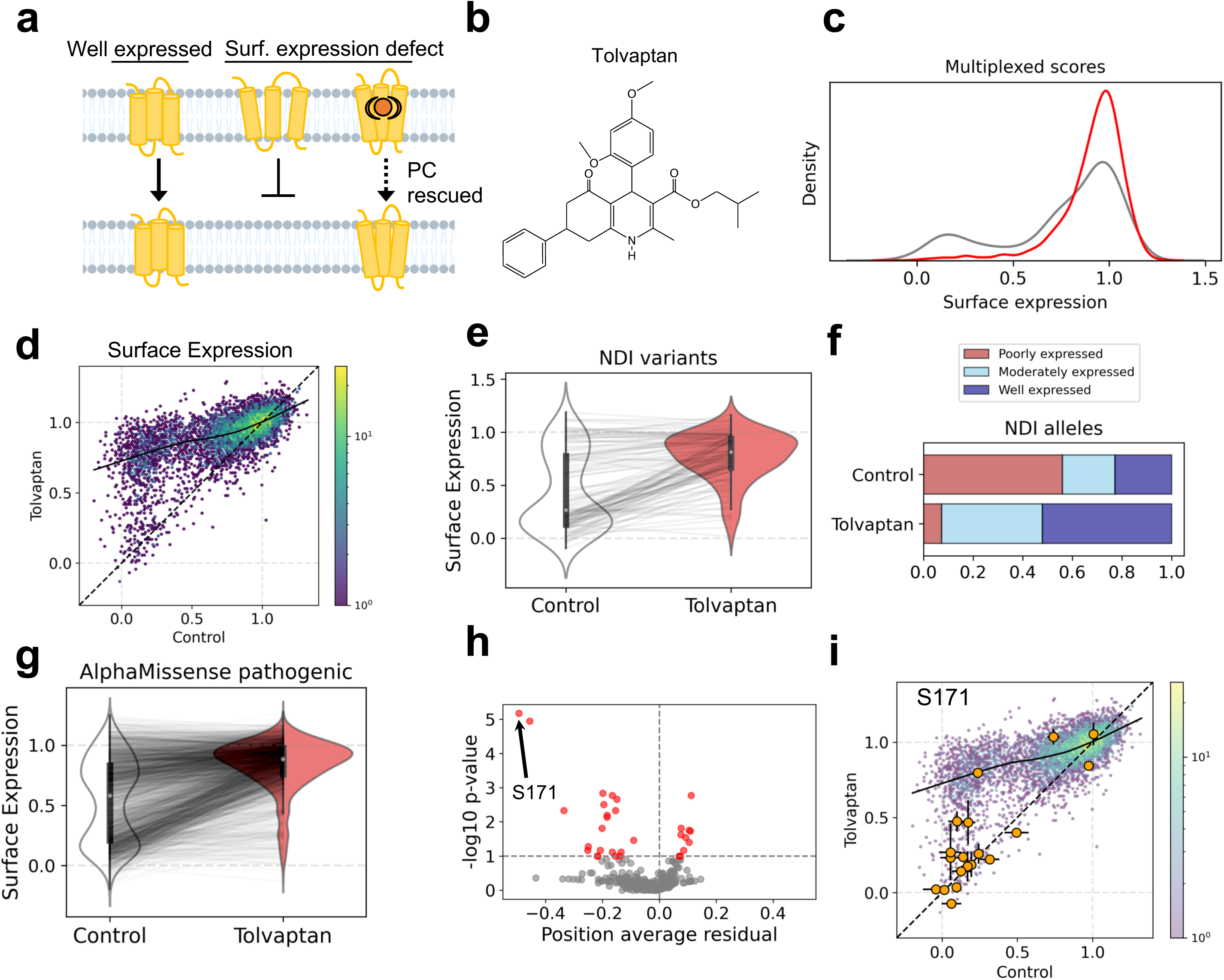
Pharmacological rescue of V2R variants. **a** Schematic showing that well-folded receptors are properly trafficked to the membrane, but folding defective variants are not properly trafficked. These experiments seek to test which variants can be rescued by pharmacological rescue. **b** Chemical structure of Tolvaptan. **c** Surface expression multiplexed score distribution of all missense variants in control (DMSO) and rescue (Tolvaptan). **d** Comparison of all missense variants in control and Tolvaptan condition; Lowess fit in solid black; x=y line in dashed black. **e** NDI variants surface expression in control compared to Tolvaptan condition. **f** Fraction of NDI variants that are poorly, moderately, or well expressed in the control and Tolvaptan conditions. **g** AlphaMissense predicted pathogenic variants surface expression in control compared to Tolvaptan condition. **h** Volcano plot comparing the average residual of all positions with the -log10 p-value. The position with the most significantly different set of residuals is highlighted (S171). **i** All missense variant effects at S171are plotted with the whole population of missense variants in the background.

Therefore, we treated cells bearing the V2R variant library with Tolvaptan for 24 hours and then profiled the population surface expression with FACS. Compared to 27°, the Tolvaptan condition has a stronger rescue effect, with more cells shifting from low to high expression (Supplemental Fig. 3b).

We sorted and sequenced the Tolvaptan rescued libraries and collected high confidence measurements for 6,759 missense and nonsense variants (91.3% of possible, data in Supplemental Table 1). Replicates were well correlated (r=0.74, Supplemental Fig. 3d) and multiplexed measurements were consistent with individual variant measurements in isogenic cell lines (r=0.85, Supplemental Fig. 3e). Multiplexed missense score distributions match the FACS distributions, and emphasize a near-complete shift of variants from low to high expression (Fig. 4c). Likewise, directly comparing surface expression levels in the presence and absence of Tolvaptan indicates nearly universal rescue (Fig. 4d).

Next, we assessed the efficacy of Tolvaptan rescue for known and predicted pathogenic variants. Tolvaptan is remarkably effective at rescuing NDI alleles: of 69 poorly expressed variants in the control condition, 60 (87.0%) are rescued to at least moderately expressed levels (Fig. 4e-f). We next evaluated effectiveness on variants predicted to be pathogenic by AlphaMissense. Strikingly, 835 of 965 poorly expressed variants (86.5%) are rescued to at least moderately expressed levels (Fig. 4g and Supplemental Fig. 3f). Tolvaptan has minor effects on NSIAD variant expression, likely because these are moderately or well-expressed already in the control condition (Supplemental Fig. 3g).

To further investigate Tolvaptan rescue, we again modeled the rescue by fitting a Lowess curve (Fig. 4d) and using residuals to this line to identify positions with more and less rescue than expected (Supplemental Table 2). More positions have specific interactions with Tolvaptan than with 27° (32 compared with 10); 21 positions are rescued less than expected, while 11 are rescued more than expected (Fig. 4h-i, FDR=0.1).

### Identifying drug binding and functional sites

Next, we sought to understand the structural features associated with rescue outlier positions. For 27° rescue, three out of ten outlier positions are glycosylation sites. Both O-glycosylation sites (S5 and T6) are rescued less than expected, but S24 is actually rescued more than expected, suggesting that temperature reduction can compensate the N-glycosylation defect (Fig. 5a, residues colored according to FDR-adjusted p value, residues outlined in yellow are rescued more than expected, those not outlined are rescued less than expected). Of the others, three are near the orthosteric site (Fig. 5a1) and two are in or near helix 8 (Fig. 5a2).

**Fig. 5.**
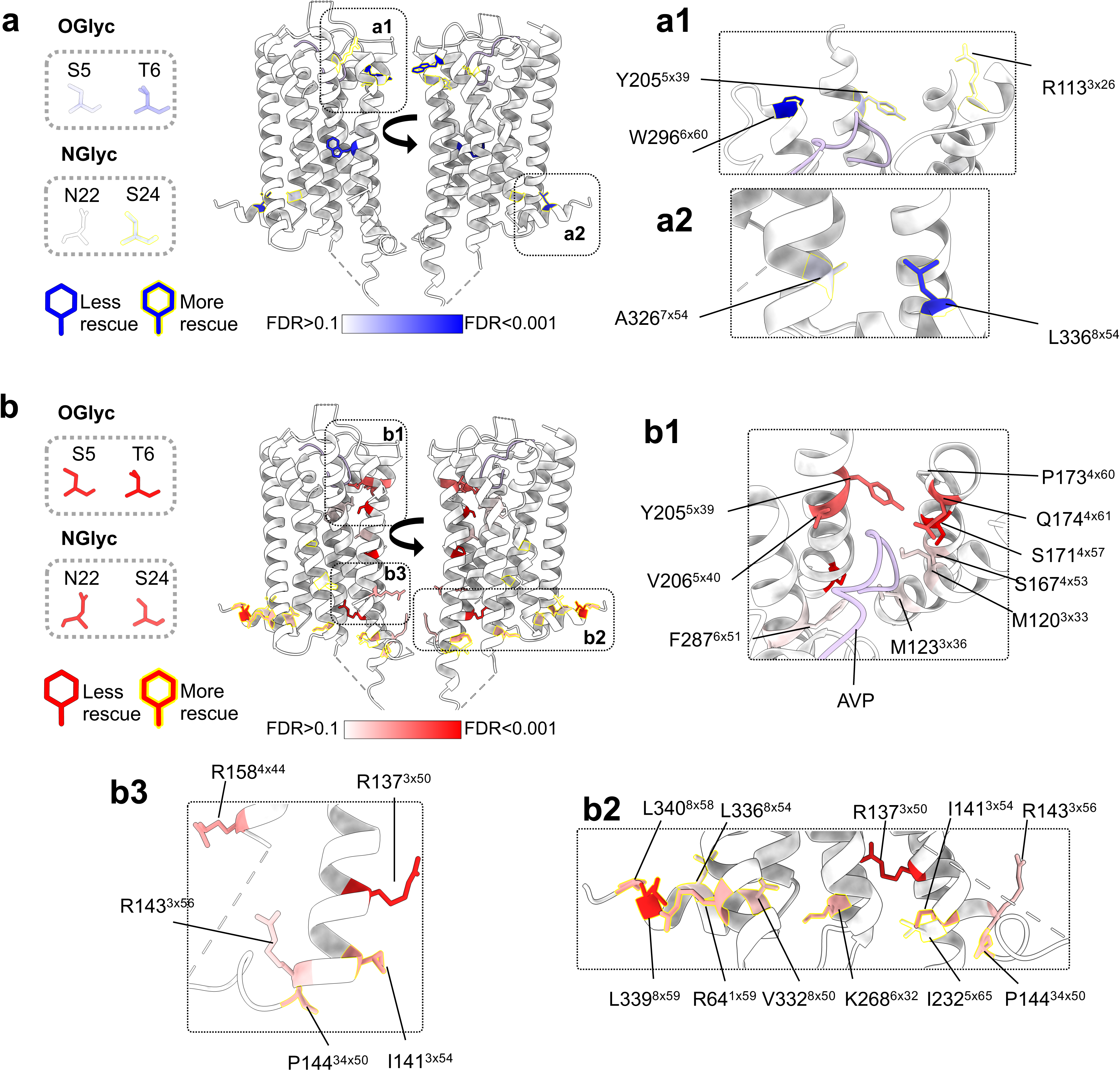
Identifying sites with more or less rescue than expected. **a** Sites that are rescued more (blue with yellow outline) or less (blue with no outline) than expected by 27°. Glycosylation sites are not resolved in the crystal structure, so these are shown individually on the left. Sites of interest outlined with dashed boxes on the structure are shown to the right. **b** Sites that are rescued more (red with yellow outline) or less (red with no outline) than expected by Tolvaptan. Sites of interest outlined with dashed boxes on the structure are shown to the right and below.

For Tolvaptan, all four glycosylation sites (S5, T6, N22, and S24) are rescued less than expected (Fig. 5b), implying that Tolvaptan cannot compensate for the lack of proper glycosylation. In addition, a cluster of residues on one side of the orthosteric binding site are rescued less well than expected (Fig. 5b1). We suspected that this could be the binding site of Tolvaptan. Indeed, a study employing molecular dynamics and site-directed mutagenesis^58^ determined that the most important residues for Tolvaptan antagonism are M123^3x36^, F178^3x36^, Y205^5x39^, V206^5x40^, and F287^6x51^, of which all but F178^3x36^ are rescued less than expected by Tolvaptan. Finally, R137^3x50^ of the highly conserved E/DRY motif and two arginines in close proximity (R158^4x44^ and R143^3x56^) are rescued less well than expected (Fig. 5b2). Mutations at R137^3x50^ cause constitutive signaling activity^46^, so this cluster of mutations might render Tolvaptan less effective by biasing the receptor to the active conformation. An examination of the sites where variants are rescued more than expected highlights a cluster of residues at the intracellular interface of the receptor (Fig. 5b3). Mutations at these sites could potentially affect signaling and/or internalization; PC stabilization of the inactive state might therefore have an exaggerated effect here.

Taken together, these results demonstrate nearly universal surface expression rescue of destabilized V2R variants with the PC Tolvaptan. The few sites where variants are consistently not rescued inform on the likely binding site of Tolvaptan as well as other functional sites of the receptor.

## Discussion

Reduced expression due to impaired fold stability is the predominant mechanism by which missense variants cause disease, including in both soluble^5,59–62^ and membrane proteins^63–66^. The destabilization conferred by most missense variants in most proteins is, however, small, suggesting that re-stabilization of a protein by a small molecule PC that binds the native state might confer sufficient free energy to rescue the effects of very diverse mutations throughout a protein’s structure. In order for this approach to be effective, though, it is necessary to prospectively classify the mechanism for all pathogenic variants in a protein, as well as their response to PC. Here we have implemented this framework and tested the ‘universal PC’ hypothesis for V2R. First, we show that over half of known loss-of-function variants are poorly expressed, and use temperature reduction to show that the vast majority of variants lose expression as a result of thermodynamic instability. Based on this, we then test the efficacy of Tolvaptan, a well-characterized antagonist and PC of V2R. Our results show that binding of a single small molecule PC can rescue the cell surface expression of nearly all variants throughout the receptor’s structure.

Previous low-throughput studies mostly tested the efficacy of PCs for a small number of variants, with the failed rescue of individual variants leading to suggestions of widespread idiosyncratic effects between PCs and variants^31–34^. More systematic efforts testing hundreds of variants in rhodopsin^14,67^ and CFTR^68^ demonstrated rescue of many variants, but with the authors also suggesting substantial region-specific differences in rescue. In contrast, we have profiled an order of magnitude more variants to demonstrate protein-wide, nearly universal PC rescue of variants.

We believe there are good empirical and theoretical reasons to expect our results to generalize to other proteins and PCs. Most proteins are marginally stable, meaning that large changes in expression result from mutations with ΔΔG of only a few kcal/mol^15^, and indeed large-scale surveys of mutational effects show most pathogenic mutations only cause small changes in fold stability^5,16,17^. It is reasonable to believe that such small changes in fold stability can be easily compensated for by small molecule binding in many proteins. Tolvaptan binding to V2R, for example, confers a free energy change of approximately -12 kcal/mol,^58^ so potentially completely offsetting the destabilizing effects of nearly all variants, provided drug concentrations are high enough.

Accordingly, small molecules may represent a much more general strategy to rescue poorly expressed variants than is commonly perceived. The law of mass action means that to rescue expression, a small molecule only needs to bind specifically to the folded conformation of a protein and with sufficient energy to shift the folding equilibrium towards the folded state (Fig. 6). There is no requirement to specifically recognize the mutated amino acid and no requirement to bind any particular region of the protein. Similar principles apply when multiple mutations are combined in a protein, where large-scale experimental analyses have revealed that changes in fold stability are nearly always additive when diverse mutations are combined in a single protein^20,69,70^. We believe, therefore, that many small molecules binding to proteins with sufficient free energy will behave as general or universal PCs.

**Fig. 6.**
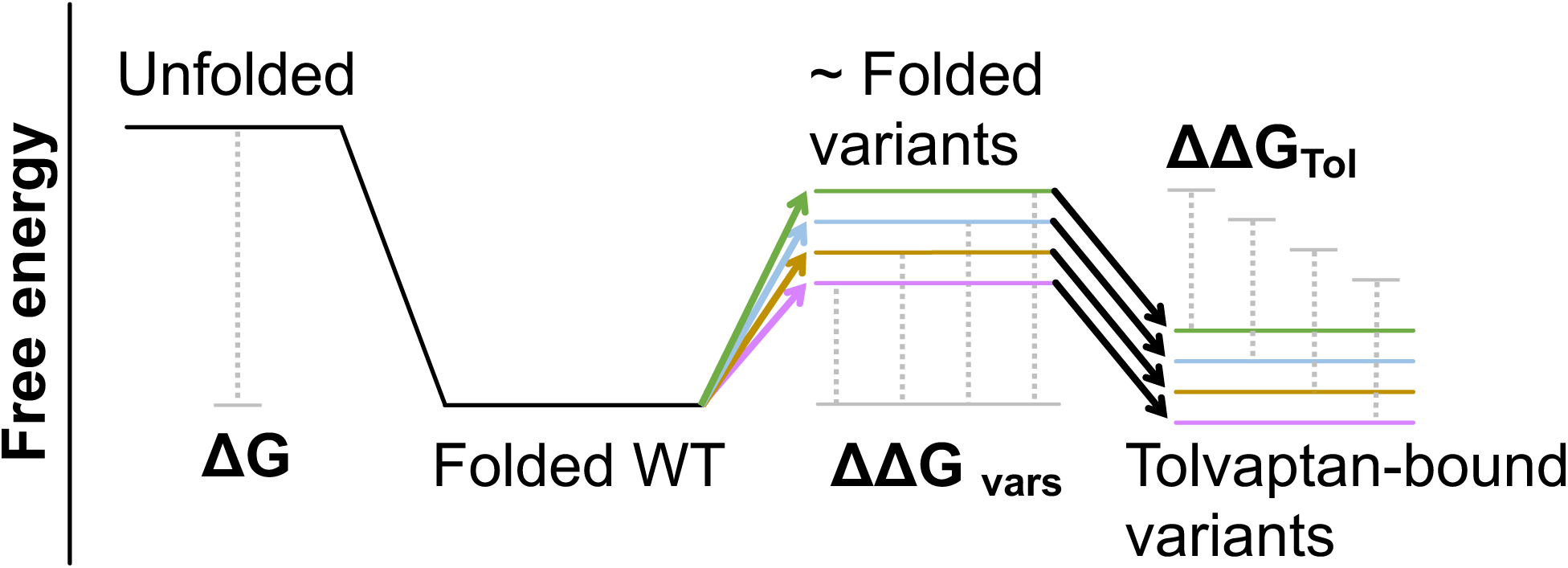
Additive model for Tolvaptan rescue. Free energy model of V2R folding, genetic variation, and rescue. We propose a model in which variants contribute different ΔΔG effects to the system, while Tolvaptan contributes a constant ΔΔG. In most cases, the ΔΔG of variants and ΔΔG of Tolvaptan combine additively.

In further support of the general, energetically additive view of PC efficacy, the small number of variants that are not rescued by Tolvaptan mostly have clear mechanistic explanations: mutations in the binding site of Tolvaptan interfere with binding of the drug and variants that affect expression by mechanisms other than reduced fold stability are not rescued, here exemplified by variants in post-translational modification sites. These variants that are not rescued by a PC can be rapidly identified by selection and sequencing experiments and so excluded from inclusion in clinical trials. We also note that this approach of identifying changes in abundance not rescued by small molecule binding is a potentially very general strategy to rapidly identify drug binding sites in proteins.

High-throughput protein abundance selection assays have now been developed for many different classes of protein^17,71,72^. As we have demonstrated here, these assays can be used to rapidly quantify the efficacy of PCs across all variants in a protein to prioritize broadly effective PCs. One limitation of our Tolvaptan rescue finding is that we have not established what fraction of rescued variants can signal at the membrane. However, there is reason to expect that a significant portion will retain signaling activity. Several studies have demonstrated detectable signaling for PC-rescued V2R^32,34,56,57^ as well as for other GPCR variants *in vitro* and *in vivo*^73,74^. Indeed a small clinical trial found that another V2R antagonist PC, SR49059, showed clinical improvement in patients, although the trial was discontinued due to off-target effects^53^.

The demonstration that a PC rescues the expression of most missense variants throughout the structure of a protein has wide-ranging implications for rare disease research. Previous experimental^5^ and computational^6,7^ approaches estimate that 40-60% of pathogenic variants are explained by loss of stability (which is in line with our findings here), suggesting a broad scope for PC therapy. Such general PCs will not have to bind to specific sites in a protein and they will not need to be tailored for each specific pathogenic variant. Rather, in accordance with the simple principle of additive free energies, any molecule that binds specifically to the folded state of a protein with sufficient free energy is a potential universal PC.

## Materials and Methods

### V2R saturation mutagenesis and barcoding

The sequence for the barcoded attB-HA-V2R construct is provided as a genbank file (https://zenodo.org/records/14216036). In order to introduce all single amino acid mutations, we used SUNi mutagenesis^35^. In short, oligos were designed to encode all amino acid mutations (with NNK or NNS degenerate codons) with flanking sequence between 18 and 40 to optimize melting temperature and presence of a 5’ GC clamp^35^. Oligos were ordered as an oPool from Integrated DNA Technologies (IDT, Supplemental Table 2). 4 μL of final product from the SUNi protocol was electroporated into 50 μL of 10β High-efficiency Electrocompetent E. coli cells (New England Biolabs (NEB)), followed by 1 hour recovery with 2 mL of SOC media. At this point, 0.5% of the recovery was plated onto an LB agar plate with ampicillin, and 99.5% was inoculated into 100 mL of LB liquid with ampicillin. Total transformant number, estimated by number of colonies on the plate, was estimated to be >200,000. Plasmid was isolated the following morning with Qiagen Plasmid Plus Midi kit.

In order to introduce the barcode construct, 5 μg of purified plasmid was digested with 250 units of ApaI restriction enzyme (NEB) for 1 hour at 37°. Then, 1.5 μL of QuickCIP (NEB) was added and the reaction was incubated at 37° for a further 30 minutes. The reaction was then run on an agarose gel and the digested fragment was isolated and column purified. The barcode (ApaI_barcode, Supplemental Table 4) was designed to have 20 degenerate nucleotides interspersed with constant AT dinucleotides: [5xN]AT[5xN]AT[5xN]AT[5xN] and flanked by sequence complementary to the sequence flanking the ApaI cut site in the attB-HA-V2R plasmid and ordered as a single stranded oligonucleotide from IDT. ApaI_barcode was made double stranded and amplified by primers amp_ApaI_barcode[F/R]. Then, the barcode was introduced into the plasmid via Gibson assembly using 60 ng of vector and 3 ng of barcode and incubated at 50° for 20 minutes. The reaction was then diluted 1:12 with water and 1.5 μL of this was electroporated into 25 μL of 10β High-efficiency Electrocompetent E. coli cells. Recovery was done in 2 mL of SOC at 37° for 45 minutes, then 0.5% was plated onto LB-agar plate with ampicillin and the other 99.5% was split between four flasks of 15 mL LB-amp liquid culture. In the morning, the number of clones on the plate was used to estimate that each flask should contain ∼25,000 transformants. Two flasks were discarded and the other two combined to arrive at an estimated library diversity of 50,000. These were further incubated another 12 hours in 100 mL of LB and finally purified with Qiagen Plasmid Plus Maxi kit.

### Long read sequencing to associate variants with barcodes

To liberate the sequence of interest from the rest of the plasmid (i.e. the barcode and V2R coding sequence), 10 μg of plasmid was digested with 15 units of KpnI-HF and 6 units of FseI (NEB), and the fragment was then run on an agarose gel and the digested fragment was isolated and column purified.

SMRTbell libraries were prepared using the SMRTbell Express Template Prep Kit 2.0 (Pacific Biosciences (PacBio)). The V2R mutagenesis libraries were multiplexed with other libraries and ran across two SMRT Cells on the Sequel IIe instrument (PacBio). The PacBio sequencing data was analyzed with alignparse^75^. Reads were quality filtered by removing any read with estimated error rate above 1x10^-4^ for the V2R coding sequence, or above 1x10^-3^ for the barcode. Then, consensus sequences were called using the alignparse.consensus.simple_mutconsensus method with default parameter settings. For further processing, barcodes with variant call support >=2 were retained, as well as barcodes with variant call support = 1 but the variant had multiple nucleotide changes compared to wild-type.

### Landing pad cell line recombination

HEK293T LLP-iCasp9-Blast Clone 12 from Matreyek et al.^38^ (hereafter referred to as “landing pad cells”) were used for library integration, expression, and screening. Cells were cultured in DMEM supplemented with 10% tetracycline-free-FBS. For recombination, 10 million landing pad cells were plated onto a T175 cell culture flask. The following day, 20 μg of library was combined with 20 μg of pCAG-NLS-Bxb1 and transfected with Lipofectamine 3000 (Thermo Fisher Scientific) per manufacturer’s instructions. After 48 hours, media was removed and replaced with doxycycline-containing media (2 μg/ml, Sigma-Aldrich). 24 hours after this, media was replaced again with media containing doxycycline as well as rimiducid (10 nM, Selleckchem). Rimiducid causes cell death in unrecombined cells and substantial cell death was apparent after 24 hours. At this point, the media was replaced with media containing only doxycycline but not rimiducid. Cells were then grown out and passaged when approaching 95% confluency, always in the presence of doxycycline.

### Cell sorting

Drug treatment was done 24 hours prior to sorting. Tolvaptan (Selleckchem, catalog number S2593) was dissolved to 10 mM in DMSO then added to cell culture media for a final concentration of 10 μM. To dissociate cells, they were first washed once with PBS, then were incubated with Trypsin-EDTA (0.05%) for 4 minutes at room temperature. Then cells were washed off the plate with media, then pelleted and resuspended in blocking buffer (1% bovine serum albumin in phosphate buffered saline). Cells were counted and 30-50M cells were transferred to a new tube. Blocking buffer was added to attain 15M cells/mL. Then, cells were incubated on a rotating wheel at 4° for 30 minutes. Following this, HA-Tag (6E2) monoclonal antibody Alexa Fluor 647 Conjugate (#3444, Cell Signaling Technologies) was added to a final concentration of 1:100, then cells were again incubated on a rotating wheel at 4° for an additional 60 minutes. At this point, cells were pelleted and supernatant removed, then resuspended in 5 mL of blocking buffer with propidium iodide (1 μg/mL).

Cells were sorted on a BD FACSaria II. Cells were first filtered by forward scattering area and side scattering area, then single cells were isolated with forward scattering width and height. BFP positive cells were filtered as unrecombined landing pad cells, and propidium iodide-positive cells were filtered as dead cells. Then, the remaining population of cells was sorted into four bins, based on Alexa Fluor 647 signal intensity, that were designed to result in a similar number of cells in each bin. For each replicate, about 10M cells were collected in total. After sorting, cells were pelleted and frozen at -80°, and processed later.

### Sequencing library preparation

DNA was isolated from cell pellets using the DNeasy Blood & Tissue Kit (Qiagen) and eluted with 150 μL of buffer EB. For each sample, 128 μL of sample was amplified in 400 μL polymerase chain reactions (PCRs) across 8 PCR tubes using Q5 High-Fidelity DNA Polymerase (NEB). Primers contained partial Illumina adapters, variable degenerate bases to promote complexity on the flowcell, and sequence complementary to the barcode flanks to amplify the barcode (ApaI_barcode_seq_[F/R]_[3-5]N, Supplemental Table 4). The PCR program was 98° for 30s; then 25 cycles of 98° for 15s, 64° for 30s, 72° for 30s; then 72° for 30s followed by 8° forever. The products from this reaction were cleaned up with NucleoSpin columns (Macherey-Nagel) and eluted in 50 μL of water. 2 μL of the eluate was then amplified in a second PCR that appended the rest of the Illumina sequencing adapter as well as index sequences (PCR2_i[5/7]). The PCR program was 98° for 30s; then 5 cycles of 98° for 15s, 64° for 30s, 72° for 30s; then 72° for 30s followed by 8° forever. The products from this second PCR were run on an agarose gel and the intended product isolated and column purified. Libraries were sequenced on an Illumina NovaSeq 6000 with 2x50 paired end reads.

### Sequencing data analysis and calculation of surface expression scores

Paired end sequencing reads in fastq format were first merged with vsearch^76^ then adapters were trimmed with cutadapt^77^. The remaining sequence corresponds to the barcode, which was compared to the full list of barcodes identified from the PacBio variant-barcode association and tallied. Then, for each variant, the frequency of that variant within each bin is calculated. This frequency is compared with the original number of cells sorted into each bin to estimate the number of cells of that genotype in each bin. Then, all barcode-variants that code for the same amino acid change are combined, and the log10 transformed geometric mean fluorescence (fluor) value of all cells sorted into each bin is combined with the estimated cell counts to arrive at the raw surface expression estimate:

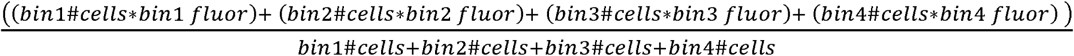

Then, these surface expression scores are normalized such that wild-type = 1 and the median of known loss-of-function variants (premature stop codons before the 300^th^ residue) = 0. Variants were retained and considered high confidence if the estimated number of cells sorted for that genotype was >=50. For the control scores, four replicates were combined-these were the controls for the rescue experiments. Namely, one condition with normal culture conditions (37° condition), two conditions with 0.1% DMSO (DMSOA and DMSOB, controlling for tolvaptan), and one condition with 1% DMSO. Since these replicates were well-correlated with each other, they were all used for deriving the control estimates.

### Modeling rescue and identifying outliers

LOWESS curves were fit to the controls vs. rescue data with the python package statsmodels.nonparametric.smoothers_lowess^78^ with these parameter values: frac=0.3, it=3, delta=0.0. After fitting the LOWESS curve, residuals were calculated for all variants as the distance in the y-axis between the curve and the variant. Then, for each position, a Mann-Whitney U test was conducted comparing the residuals of all variants at that position with control surface expression below 0.85 with all other variants with control surface expression below 0.85. This threshold was intended to only include variants that actually had the potential to be rescued. To control the false discovery rate, we then used statsmodels.stats.multitest.multipletests, method=”fdr_bh”, alpha=0.1.

### Clinical variant curation

gnomAD data was version v2.1.1, controls only, and was obtained on February 8, 2024. Clinvar variants were obtained on February 9, 2024 and filtered for missense variants. HGMD variants were obtained on February 8, 2024 and filtered for missense variants.

### Computational predictors

ESM1b was installed and ran as described (https://github.com/ntranoslab/esm-variants), using the esm_score_missense_mutations.py function. AlphaMissense, EVE, and RaSP scores were available as precomputed scores that we downloaded from the respective servers. ThermoMPNN scores were computed using the Google Colab server.

## Acknowledgments

This work was funded by a European Research Council Advanced Grant (883742), the Spanish Ministry of Science and Innovation (LCF/PR/HR21/52410004, EMBL Partnership, Severo Ochoa Centre of Excellence), the Bettencourt Schueller Foundation, the AXA Research Fund, Agencia de Gestio d’Ajuts Universitaris i de Recerca (AGAUR, 2017 SGR 1322) and the CERCA Program/Generalitat de Catalunya. T.L.M. was funded by an EMBO fellowship (ALTF 113-2021). We thank all members of the Lehner Lab and Jana Selent and Tomasz Stepniewski for helpful discussions. We thank the CRG/UPF Flow Cytometry Unit for assistance with the sorting experiments.

## Data and code availability

Files needed to reproduce analyses can be found at zenodo (https://zenodo.org/records/14216036). Custom code to reproduce analyses can be found at github (https://github.com/lehner-lab/V2R_surfexp_rescue). Raw sequencing reads can be found at Sequence Read Archive (accession number PRJNA1190688).

