## Supplementary figures for "A pharmacological chaperone stabilizer rescues the expression of the vast majority of pathogenic variants in a G protein-coupled receptor"

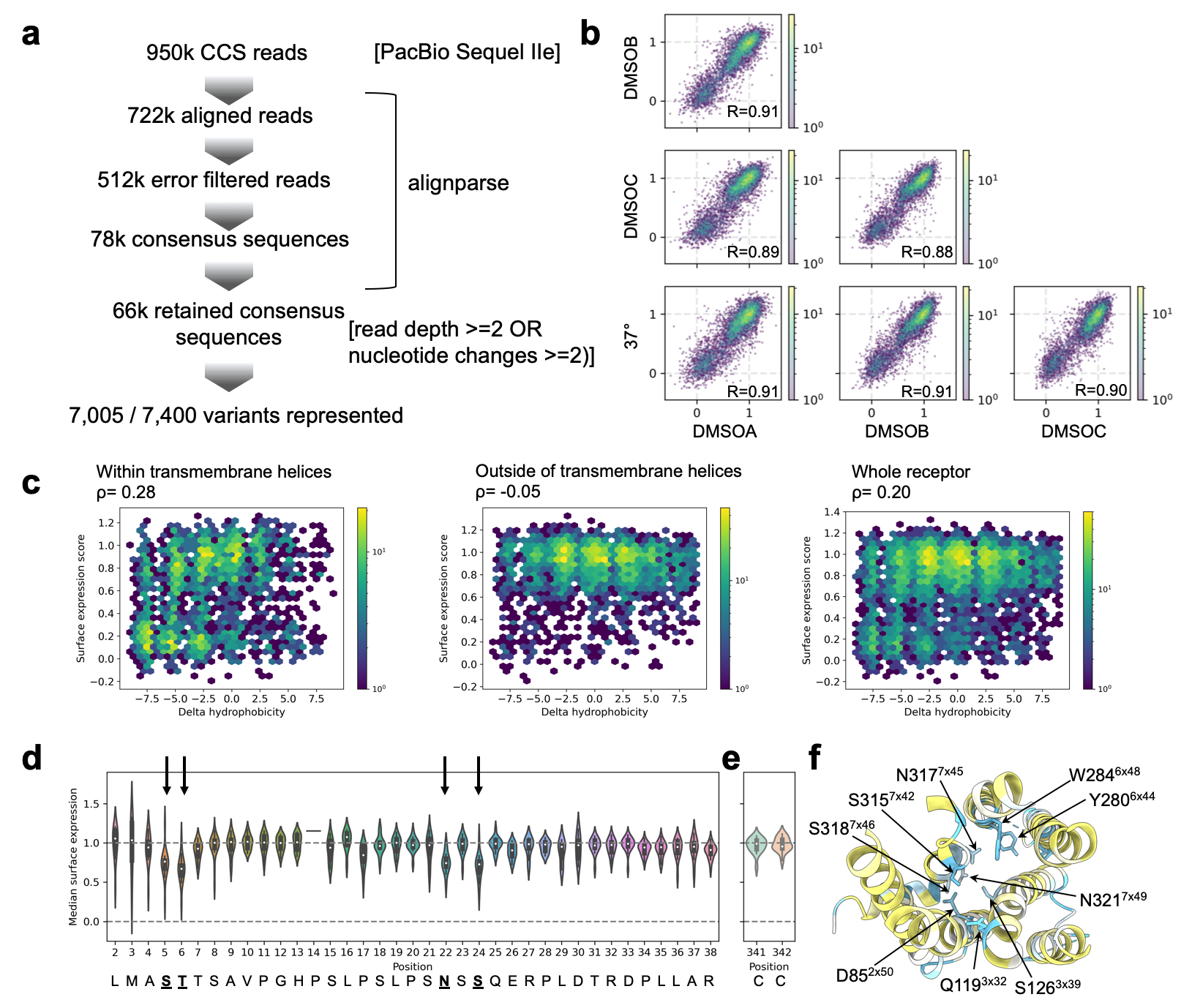

**Supplementary Fig. 1. Extended analyses of primary V2R surface expression data.**

**a** Overview of data processing for barcode-variant association with PacBio long read sequencing. **b** Replicate correlations for the four control conditions. 37° was culture media without DMSO, DMSOA and DMSOB were culture media with 0.1% DMSO, DMSOC was culture media with 1% DMSO. Because the conditions are all well correlated, they were all considered control and combined for the control estimates. **c** Hexbin plots comparing the change in hydrophobicity with surface expression score, separated by positions that are in transmembrane domains, outside of transmembrane domains, or the whole receptor considered together. **d** Violin plot of variant effects in the N-terminus of V2R. Highlighted with arrows are the putative O- and N-glycosylation sites (5 and 6, 22 and 24, respectively). **e** Variant effects at the sites of palmitoylation. **f** Topdown view of V2R, with residues colored by their preference for hydrophobicity. Color scale as in Fig. 1h. Highlighted with arrows are residues in the core of the receptor where hydrophilic amino acids are preferred.

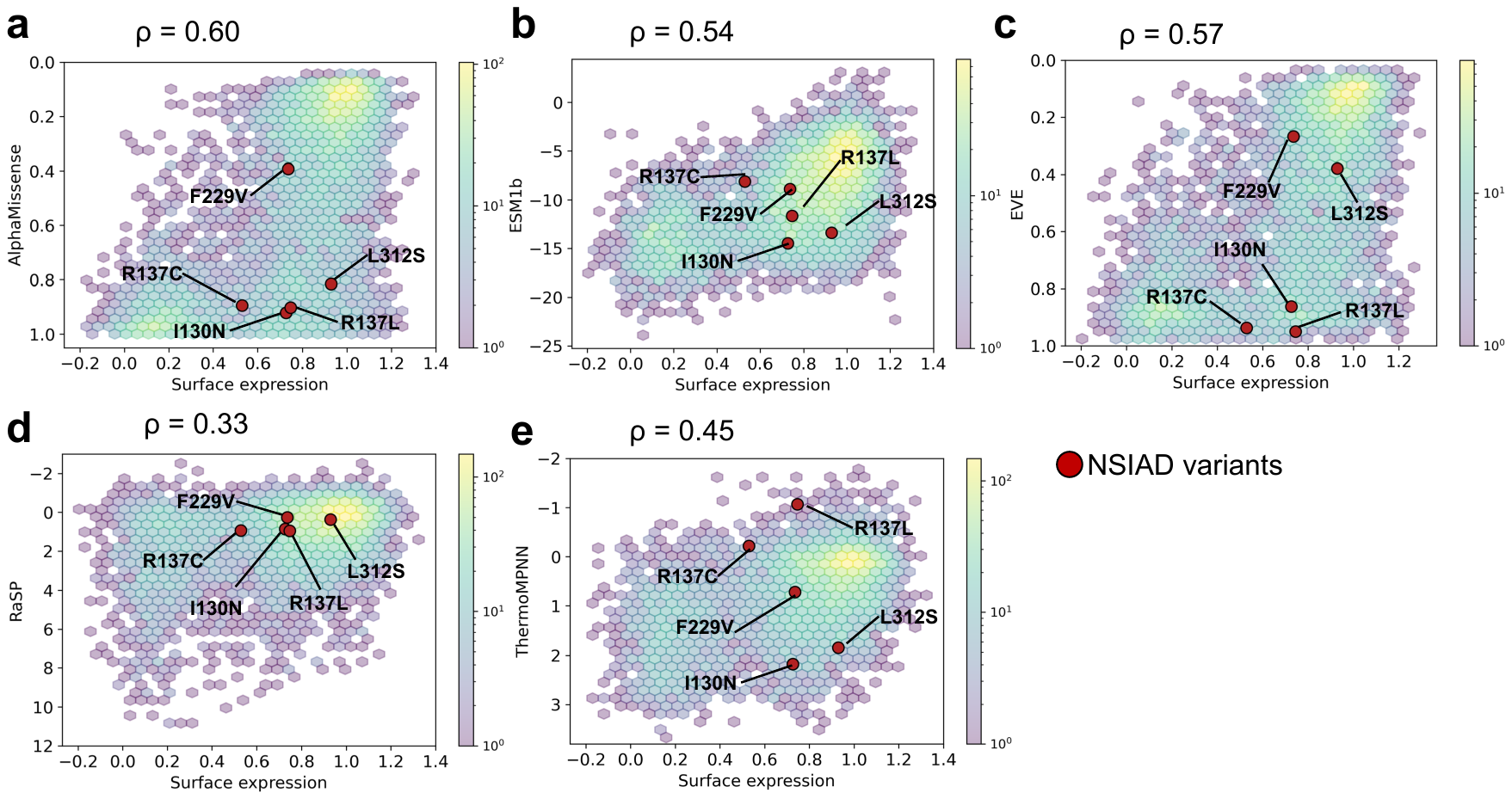

**Supplementary Fig. 2. Comparison of computational VEPs with empirical surface expression scores, highlighting NSIAD scores.**

**a** Hexbin plot comparing surface expression with AlphaMissense scores for all missense variants. NSIAD scores are highlighted. **b** Hexbin plot comparing surface expression with ESM1b scores for all missense variants. NSIAD scores are highlighted. **c** Hexbin plot comparing surface expression with EVE scores for all missense variants. NSIAD scores are highlighted. **d** Hexbin plot comparing surface expression with RaSP scores for all missense variants. NSIAD scores are highlighted. **e** Hexbin plot comparing surface expression with ThermoMPNN scores for all missense variants. NSIAD scores are highlighted.

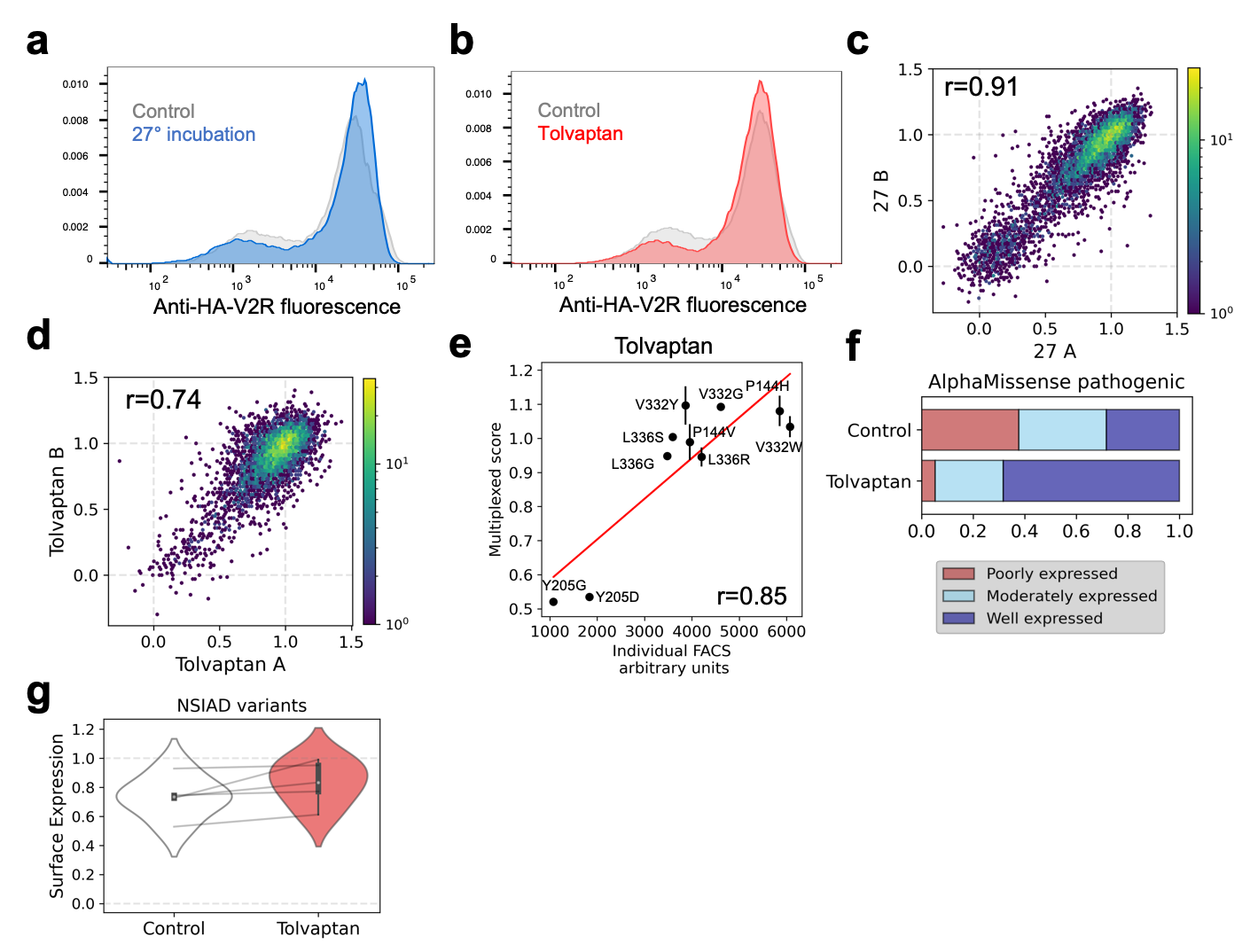

**Supplementary Fig. 3. Extended analysis of rescue experiments.**

**a** FACS data for V2R library in the control versus reduced temperature (27°) condition. **b** FACS data for V2R library in the control versus Tolvaptan condition. **c** Replicate correlations for 27° experiment. **d** Replicate correlations for Tolvaptan experiment. **e** Comparison of ten variants measured in multiplex or individually, in the presence of Tolvaptan. **f** Percent of AlphaMissense predicted pathogenic variants that are poorly, moderately, or well expressed in control and Tolvaptan conditions. **g** Violinplot comparing the surface expression scores of NSIAD variants in the control compared with the Tolvaptan condition.
